# Analysis of genetic variation in *Macadamia* shows evidence of extensive reticulate evolution

**DOI:** 10.1101/2024.02.15.580603

**Authors:** Sachini Lakmini Manatunga, Agnelo Furtado, Bruce Topp, Mobashwer Alam, Patrick J. Mason, Ardashir Kharabian-Masouleh, Robert J Henry

## Abstract

The genus *Macadamia* in the Proteaceae family includes four species. To explore diversity in macadamia genetic resources, a total of 166 wild genotypes, representing all four species, were sequenced. The four species were clearly distinguished as four separate clades in a phylogenetic analysis of nuclear gene CDS. *M. integrifolia* and *M. tetraphylla* formed a clade that had diverged from a clade including *M. ternifolia* and *M. jansenii.* The greatest diversity in nuclear and chloroplast genomes was found in the more widely distributed *M. integrifolia* while the rare *M. jansenii* showed little diversity. The chloroplast phylogeny revealed a much more complex evolutionary history. Multiple chloroplast capture events have resulted in chloroplast genome clades including genotypes from different species. This suggests extensive reticulate evolution in *Macadamia* despite the emergence of the four distinct species that are supported by the analysis of their nuclear genomes. The chloroplast genomes showed strong associations with geographical distribution reflecting limited maternal gene movement in these species having large seeds. The nuclear genomes showed lesser geographical differences, probably reflecting longer distance movement of pollen. This improved understanding of the distribution of diversity in *Macadamia* will aid conservation of these rare species now found in highly fragmented rainforest remnants.

**Highlights:** Whole genome sequencing of population of the four species in the Macadamia genus allowed phylogenetic relationships to be determined and revealed significant reticulate evolution.

## 1. Introduction

Phylogenetics, the study of evolutionary relationship among organisms, has become a powerful tool in a variety of biological disciplines (Liu *et al*., 2022). Over the last few decades, substantial effort has been made to understand the phylogenetic associations among angiosperms through the application of DNA sequence data (Jansen *et al*., 2006). Next Generation Sequencing (NGS) has brought a transformation in sequence analysis by making it more affordable and increasing access to complete chloroplast and nuclear genome data (Zhou *et al*., 2021). There has been an emphasis by breeders to exploit wild germplasm, but a significant portion remains underutilized (Zhang and Batley, 2020). The lack of genomic data on wild germplasm is one barrier to effectively use it in plant breeding efforts and as a result the integration of genes from wild germplasm into cultivated varieties has been limited. Recent advancements in genomics and bioinformatics have provided opportunities to unlock the hidden potential within wild germplasm (Tanksley, 1997) by extending this technology to less studied plant species. This has opened new avenues to incorporate materials from wild germplasm (Zhang and Batley, 2020).

Macadamia is an evergreen perennial rainforest tree of the family Proteaceae and is indigenous to Australia (Hardner *et al*., 2009; O’Connor *et al*., 2019; Walton, 2011). The genus *Macadamia* is the only angiosperm that has been domesticated as a large scale commercial food crop in Australia (Aradhya *et al*., 1998). In accordance with the present classification of Proteaceae (Mast *et al*., 2008) the genus *Macadamia* has been classifying into four species, namely; *M. integrifolia* (Maiden & Betche), *M. tetraphylla* (L.A.S. Johnson), *M. ternifolia* (F. Muell) and *M. jansenii* 9C.L. Gross & P.H. Weston) using molecular and morphological data, while many species previously classified as *Macadamia* have been transferred to other genera. Among the four species, *M. integrifolia* has the widest natural distribution, extending from south-east Queensland to the New South Wales border. Two overlapping distributions lead to natural hybridization between *M. integrifolia* and *M. tetraphylla* and between *M. integrifolia* and *M. ternifolia* (O’Connor *et al*., 2019; Topp *et al*., 2019). *M. tetraphylla* is mostly distributed in New South Wales while *M. ternifolia* is distributed north of Brisbane extending from the Samford Valley to Gympie (Topp *et al*., 2019). *M. jansenii* is the most geographically isolated species and is found only in Bulburin National Park north of Bundaberg, 180 km from the closest *M. integrifolia* population (Mai *et al*., 2020; Topp *et al*., 2019). Of the genus *Macadamia* display diversity in several morphological characteristics. These include the number of leaves per whorl, mature leaf size and shape, colour of new leaves, presence of petiole, leaf margin serration, as well as differences in floral and fruit morphology (Peace, 2005). These characteristics are considered for differentiating *Macadamia* species. However, some of these characteristics, such as leaf serration, can overlap across species and can be observed only a certain stage (i.e., juvenile or adult) of the lie cycle of some species. On the other hand, traits like nut and leaf size, can vary within species depending upon environment and may not always be useful in distinguishing between the species (Hardner *et al*., 2009; Peace, 2005). Genomic information on the representative accessions of these four species can be instrumental in understanding the diversity and species distribution of *Macadamia*.

Although genomic investigation in macadamia commenced a decade ago, only a few studies have been conducted till to date. Several types of polymorphic molecular markers have been used to assess the genetic diversity in *Macadamia* (Ahmad Termizi *et al*., 2016; Alam *et al*., 2018; Mai *et al*., 2020; Mast *et al*., 2008; O’Connor *et al*., 2019; Peace, 2005). However, few studies have been employed to characterize the genetic makeup of wild germplasm (Mai *et al*., 2020). In 2005, Peace *et al*. studied a large number of wild germplasm accessions using low throughput RAF (randomly amplified DNA fingerprinting) and RAMiFi (randomly amplified microsatellite fingerprinting) markers (Peace, 2005). Another study by Mast *et al*. (2008) investigated the relationships between the four *Macadamia* species and their closely related wild relatives. They examined chloroplast DNA regions, such as matK, atpB and ndhF and nuclear DNA genomic regions such as waxy loci 1 and 2 and PHYA. By analysing these markers their aim was to gain insights into the complex relationships, within the *Macadamia* genus and its wild relatives (Mast *et al*., 2008). However, these markers gave low genome coverage and provided poor marker density (Alam *et al*., 2018; Nock *et al*., 2020). Ahmad *et al*. (2016) analysed individuals from wild *M. integrifolia* population using 516 Single Nucleotide Polymorphisms (SNPs) and reported the unique chlorotypes for each of 12 samples (Ahmad Termizi *et al*., 2016). Furthermore a recent study (Mai *et al*., 2020) examined the genetic relationships among 302 accessions of wild germplasm using 2872 SNP and 8415 silico DArT markers and identified the species status of 94 unknown wild accessions. Although these studies examined the phylogenetic relationships among wild macadamia accessions, no previous study has resolved the phylogeny of the four *Macadamia* species.

In *Macadamia*, as in other plants uniparentally inherited chloroplast DNA has been used to infer the phylogenetic patterns. However many studies have documented the occurrence of reticulate evolution of chloroplast in other plant species (Nge *et al*., 2021). The phenomenon of reticulate evolution may result in the replacement of chloroplast genomes of one species with another (Kawabe *et al*., 2018) due to hybridization events. In many plants, reticulate evolution has caused a discordance between the molecular data derived from the chloroplast and nuclear genome (Nge *et al*., 2021) resulting in conflicting topologies for phylogenetic trees (Rieseberg, 1991). Therefore, reticulate evolution can have an impact on phylogenetic analyses that rely only on the chloroplast genomes or its genes (Kawabe *et al*., 2018). However, studies to date have not explored phylogenetic relationships in *Macadamia* based on both nuclear and complete chloroplast genomes.

Here we focused on uncovering the diversity and relationships in wild *Macadamia* populations by using both chloroplast genomes and nuclear gene coding sequences. This is the first report of whole genome sequence data for a large *Macadamia* population. To support improved conservation and utilization of the wild genetic resources we sequenced whole chloroplast and nuclear genomes to better understand diversity within and relationships between species and populations of *Macadamia* in Australia.

## 2. Materials and Methods

### 2.1. Plant materials, DNA extraction and sequencing

A total of 166 wild macadamia accessions representing all four species (*M. integrifolia* (n =49), *M. tetraphylla* (n=56), *M. ternifolia* (n=23), *M. jansenii* (n= 23)) and one related rainforest species from the Proteaceae, *Lasjia whelanii* were selected for sequencing. Within the macadamia populations, 161 wild accessions, which were collected previously from multiple locations across the natural distribution of the four species, were grown and maintained at Nambour arboretum and *ex situ* germplasm centres at Nambour and Tiaro in Queensland and Alstonville in NSW (Hardner *et al*., 2004) and five from a private collection at Limpinwood, NSW (Supplementary Table 1) in Australia. Fully expanded young macadamia leaves, of accession within these *ex situ* collection sites, were collected in perforated labelled cellophane bags and immediately placed under dry ice until stored in a in a -80 ^0^C freezer at The University of Queensland, Brisbane, Australia.

Frozen leaves were coarse pulverized under liquid nitrogen using a mortar and pestle and further fine pulverized under cryogenic condition using a Qiagen tissue lyser (MM400, Retsch, Germany). A modified version of the cetyltrimethylammonium bromide (CTAB) DNA extraction protocol described by Furtado *et al*. (2014) (Furtado, 2014) was used to extract genomic DNA. The quality and quantity of the DNA samples were evaluated using a Nanodrop spectrophotometer (Nanodrop Technologies, Wilmington, DE, USA) by recording the absorbance ratios at 260/280 and 260/230 followed by running a 0.7% agarose gel with SYBR safe staining (Thermo Fisher Scientific). Whole genome short read sequencing was undertaken by BGI Hong Kong. A PCR-free library was generated and sequencing at 150bp paired end reads was undertaken on the DNBSEQ-G400 sequencing platform from MGI (MGI Tech Co., Ltd, Shenzhen, China) at an expected data yield/sample of at least 25X genome size (800 Mb genome).

### 2.2. Chloroplast genome assembly and annotation

All sequence data were analysed in CLC Genomics Workbench 23.0.05 (CLC-GWB, CLC-Bio, QIAGEN, Denmark, http://www.clcbio.com) using the short-read pipeline. Quality control (QC) was performed for all short-read data. Reads were trimmed using a quality score limit of 0.01 with default parameters (More than 98% of the resulted trimmed reads had a Phred score >25). Subset of quality trimmed short reads (2 - 13 GB) were used for chloroplast genome assembly. All chloroplast genomes were assembled using GetOrganelle toolkit (Jin *et al*., 2020) exploiting SPAdes v.3.15.3, Bowtie2 v.2.4.5 and Blast v.2.11.0 as dependencies. The correct configuration of the chloroplast genome was selected with respect to the *M. integrifolia* (Reference sequence: NC_ 025288.1) (Nock *et al*., 2014) available at National Centre for Biotechnology Information (NCBI) ((http://www.ncbi.nlm.nih.gov/) using clone manager software (Sci Ed, USA).

The chloroplast genomes were annotated using the GeSeq online tool (https://chlorobox.mpimp-golm.mpg.de/geseq.html) with *M. integrifolia* (Reference sequence: NC_ 025288.1) as the reference genome (Tillich *et al*., 2017). Chloroplast genomes were annotated with following settings: Annotation options: Annotate plastid inverted repeat (IR), Annotated plastid trans spliced rps12, Annotation support: Support annotation by Chloe, Annotation revision: Keep best annotation only, BLAT search – protein search identity: 25, rRNA, tRNA, DNA search identity: 85, HMMER profile: CDS+rRNA, ARAGORN v1.2.38-Genetic code: Bacterial/plant chloroplast, Max intron length: 3000, tRNAscan-se v2.0.7-sequence source: organellar tRNAs, MPI-MP reference set: chloroplast land plants (CDS + rRNA) and Chloe v0.1.0-annotate-CDS + tRNA + rRNA. All genomes were imported to Geneious 2023.2.1 software (Biomatters Ltd, USA) to determine number of genes, coding sequence (CDS), transfer RNAs (tRNAs) and ribosomal RNAs(rRNAs) in each sample.

### 2.3. Concatenated nuclear gene CDS sequences

A previously published annotated sequences of the nuclear genome of *M. integrifolia* (Nock *et al*., 2020) were selected reference sequences to generate accession-specific consensus coding sequences (CDS) of nuclear genes. CDS of *M. integrifolia* (GCF 013358625.1) were downloaded and imported to CLC-GWB to generate a local Blast database. CDS of one hundred and six *Arabidopsis thaliana* single copy genes identified by Li *et al.,* (2017) (Li *et al*., 2017) was subjected to tblastn against the *M. integrifolia* CDS database. We selected 56 tblast hits with a single *M. integrifolia* CDS matching *a A. thaliana* CDS as these hits represented single copy genes in the *M. integrifolia* genome. From these selected single hits, corresponding *M. integrifolia* CDS sequences were extracted and used as a reference to extract consensus sequences from each of the macadamia accessions and from *L. whelanii*. BLAST analysis using the 56 extracted *M. integrifolia* CDS and the CDS sequences of *L. whelanii* as a database, resulted in the selection of 53 *L. whelanii* CDS as single hits which represented single copy genes in *L. whelanii*, Corresponding (same as selected) 53 CDS sequences from *M. integrifolia* and from *L. whelanii* were selected for further analysis. The 53 CDS from *M. integrifolia* (Supplementary Table 3) were used as reference sequences in the read mapping approach to generate corresponding CDS consensus sequences for each of the macadamia accessions. Essentially, short reads trimmed data of each macadamia accession were mapped separately to each of the 53 selected *M. integrifolia* CDS sequences to extract consensus sequences. Finally, consensus CDS sequences, extracted for each macadamia accessions, were concatenated in the same sequential order to obtain the final nuclear gene CDS sequence.

### 2.4. Phylogenetic evaluation

Phylogenetic evaluation of macadamia was conducted by utilizing complete chloroplast genome sequences and single copy concatenated nuclear gene CDS sequences. For phylogenetic analysis we selected 138 wild macadamia accessions representing all four species. To maintain precision and clarity within our analysis, we exclude accessions from planted wild germplasm, accessions known to be natural hybrids, admixtures, accessions of unknown origin, and unidentified macadamia accessions (Supplementary table 4). The phylogenetic trees were also generated for four species: *M. integrifolia* (n =44), *M. tetraphylla* (n=49), *M. ternifolia* (n=22), *M. jansenii* (n= 23) separately based on chloroplast genomic data and single copy nuclear gene sequences. *L. whelanii* was used as an outgroup. All selected sequences were aligned along with the outgroup using MAFFT alignment with default parameters in Geneious 2023.2.1 software (Biomatters Ltd, USA).

#### 2.4.1. Chloroplast phylogenetic analysis

Chloroplast genomes are widely used in plant phylogenetic analysis (Yanfei *et al*., 2023). Therefore, to investigate the relationships in genus *Macadamia,* phylogenetic trees were constructed using complete chloroplast genome sequences. To better determine the relationships within the *M*acadamia species, we first constructed ML trees and Bayesian trees for all four species separately using *L. whelanii* as an outgroup.

Chloroplast phylogenetic reconstructions were performed using PAUP*v 4.0 software (Swofford, 2003) with Maximum Likelihood (ML) method and MrBayes v. 3.2 software (Ronquist *et al*., 2012) in Geneious for Bayesian inference (BI) method. For PAUP* trees, the Akaike Information Criterion (AIC) in jModel test was performed in Cyberinfrastructure for Phylogenetic Research (CIPRES) Science Gateway (https://www.phylo.org/) to find out, the best fitting nucleotide substitution model (Miller, 2010). ML analysis was performed with 1000 bootstrap replicates. The GTR + Gamma was used in BI analysis. The chloroplast ML trees for *M. integrifolia* and *M. tetraphylla* population were generated by TPM1uf+I+G model and chloroplast ML trees for *M. ternifolia* was generated by TVM+G model. The topological structures of trees were evaluated based on bootstrap support and Bayesian posterior probabilities. Interactive Tree Of Life (iTOL) v.5 tool (Letunic and Bork, 2021) (https://itol.embl.de/about.cgi) was used to visualize the phylogenies.

To further evaluate the phylogenetic relationship between the four species of macadamia, we constructed Bayesian tree methods (GTR + Gamma model) by taking 138 complete macadamia chloroplast genomes using *L. whelanii* as an outgroup.

#### 2.4.2. Nuclear gene phylogenetic analysis

Concatenated 53 nuclear gene CDS sequences of *Macadamia* species and *L. whelanii* were used to evaluate the phylogenetic relationships in Genus *Macadamia*. The nuclear gene phylogenetic trees were generated by using Randomized Axelerated Maximum Likelihood (RAxML) version 8 (Stamatakis, 2014) with Maximum Likelihood (ML) method and BI method in MrBayes v. 3.2 software (Ronquist *et al*., 2012) in Geneious 2023.2.1 software (Biomatters Ltd, USA). ML trees were analysed using GTR + GAMMA nucleotide model with 1000 bootstrap replicates. The BI trees were analysed using GTR + GAMMA model. The phylogenetic trees were visualized with the iTOL v.5 tool (Letunic and Bork, 2021). The topological structure of trees produced by RAxML and MrBayes software were compared to identify discrepancies between them.

### 2.5. Assessment of phylogeography

Geographical maps of origin were created for all four macadamia species based on the chloroplast phylogenetic clade separation. Geographic ranges were mapped using Esri National Geographic in QGIS Geographic Information System (Version 3.32.1-Lima) (http://qgis.osgeo.org). Geographical coordinates of each accession are listed in Supplementary table 4.

## 3. Results

### 3.1. Characterization of chloroplast genomes

Following paired end sequencing (150 bp), a total of 140,789,058 – 260,592,896 reads were obtained for 166 *Macadamia* accessions (Supplementary table 2) with sequence depth between 18.23X – 50X. Sequence depth of trimmed paired end reads at quality score limit of 0.01 range between 16.47X – 43.98 X. Two assembled sequences were obtained from GetOrganelle analysis indicating the presence of two structural haplotypes of chloroplast genome that occurs in plants related to the orientation of the single copy region. The correct configuration of the chloroplast genome was selected with respect to *M. integrifolia* (Reference sequence: NC_ 025288.1). Complete chloroplast genome sizes analysed in this study are shown in Supplementary table 5. Complete circular chloroplast genomes were obtained for all genotypes, ranging in size from 159,195 bp – 159,734 bp. The smallest chloroplast genome was identified for three *M. tetraphylla* accessions (Mac_297, Mac_338, Mac_345) and two wild macadamia trees of uncertain species (Mac_047, Mac_329) while the largest was observed for two *M. integrifolia* accessions (Mac_029 & Mac_262). Chloroplast genome sizes of *M. integrifolia* ranged from 159,458bp – 159,734bp, *M. ternifolia* from 159,463 – 159,508 bp, *M. tetraphylla* from 159,195bp – 159,598 bp, and all *M. jansenii* were 159,524 bp in length except for MacP_16 (159,526 bp). The result of this study also revealed that out of 23 *M. jansenii* accessions, 22 accessions had identical chloroplast genomes.

All macadamia chloroplast genomes showed quadripartite structure of angiosperm including a large single copy (LSC), a small single copy (SSC) and two identical Inverted Repeats (IRa & IRb) (Figure 1). Gene annotation showed 116 full length genes, 81 coding sequences, 4 ribosomal RNA (rRNA) and 31 transfer RNA (tRNA). Among these 116 genes, 60 genes involved in protein synthesis and DNA replication (Genes responsible for ribosomal RNAs, transfer RNAs, Large subunit of ribosome, Small subunit of ribosome and DNA dependent RNA polymerase), 46 in photosynthesis (Genes responsible for subunits of photosystem I, subunits of photosystem II, subunits of ATP synthase, subunits of NADH dehydrogenase, large subunit of rubisco, subunits of cytochrome complex), 6 in different other functions (Genes responsible for inner membrane protein, cytochrome synthesis gene, acetyl-CoA-carboxylase, maturase, ATP-dependent protease and translational initiation factor) and 4 in unknown function genes (Supplementary table 6).

**Figure 1:**
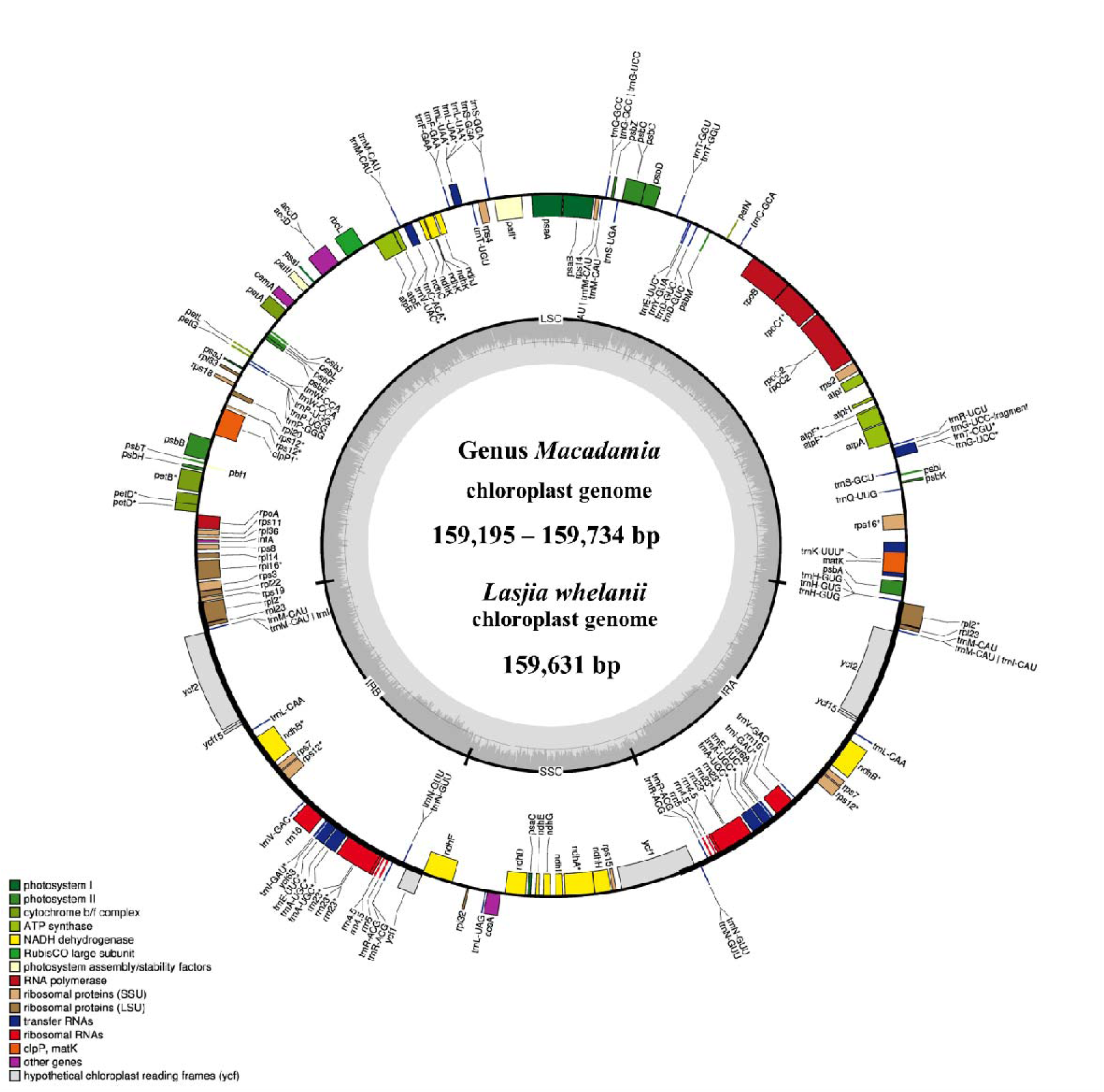
(a) The circular map of four *Macadamia* species. (b) The circular map of *Lasjia whelanii*. Genes inside the circle are transcribed in the clockwise direction whereas the genes outside the circle are transcribed in the counterclockwise direction. Genes belonging to different functional groups are colour coded. Gray area in the inner circle indicates the GC content of the chloroplast genome. The four regions of a chloroplast genome are also indicated in the inner circle: the two inverted repeat regions (IRA and IRB) are separated by small (SSC) and large (LSC) single copy regions.

The *L. whelanii* chloroplast genome possessed the standard quadripartite structure, containing two inverted repeats (18,824 bp), LSC region (87,911 bp) and SSC region (26,448 bp). (Figure 1). Genome size was recorded as 159,631 bp. Plastome of *L. whelanii* contained no significant difference in related to genes, protein coding genes, rRNA and tRNA. Overall, the GC content of the chloroplast genome was recorded as 38 %.

### 3.2. Chloroplast phylogeny and geographical analysis of *Macadamia* species

The multiple chloroplast genome alignment of 44 *M. integrifolia* accessions together with the outgroup *L. whelanii* was 161,281 bp in length with 99.6% identical sites. The tree topologies of both were similar (Figure 2a), and most nodes were supported by high bootstrap support (BS) (> 95%) and Bayesian posterior probabilities (PP) (>0.95). However, some internal nodes tended to have low BS, indicating incomplete lineage sorting. The phylogenetic tree construction revealed Mac_ 232 (Correspond to population site 90) cluster separately from rest of the 43 accessions. Remaining accessions were differentiated into two main clades and further differentiated into sub clades. Clade II contained accessions from the northern distribution of *M. integrifolia*: Mac_231, Mac_262, Mac_029, Mac_265 and Mac_033 from Gundiah/Mount Bauple region (Corresponded to population site 1, 2, 2, 3 and 3 respectively) (Figure 2b) and Mac_052, Mac_091, Mac_340, Mac_248, Mac_266 from Gympie region (Correspond to population site 9, 55, 56, 57 and 57 respectively). Clade III included a total of nine accessions, in which seven were from Caboolture region: Mac_250, Mac_312, Mac_246, Mac_045, Mac_080, Mac_026, Mac_143 (Correspond to population site 20, 21, 21, 71, 76, 76 and 77 respectively) and one from Nambour region: Mac_044 (Correspond to population site 101). Interestingly, Mac_059 from population site 57, did not cluster with other two accessions (Mac_ 248 and Mac_266) from population site 57 (Gympie region) in Clade I. Clade IV contained 24 accessions which were collected from the region south of Brisbane except for Mac_251, Mac_089, Mac_139, Mac_189 (Correspond to population site 3, 90, 90 and 90 respectively). This observation verified that the accessions having same geographical origin tend to form distinct clusters among themselves. However, it is noteworthy that certain accessions from these localities exhibit a tendency to associate or cluster with accessions sourced from different geographical areas. The result of this study also revealed that Mac_251, Mac_089, Mac_139, Mac_189 (Correspond to population site 3, 90, 90 and 90 respectively) are likely to be planted trees. Mac_312, Mac_246 and Mac_306, Mac_228 are biological replicates and confirmed by clustering together. Additionally, short branch length of the tree suggested that Mac_232 had less divergence while Mac_026, Mac_044, Mac_080 and Mac_143 from Clade III were more diverged accessions.

**Figure 2:**
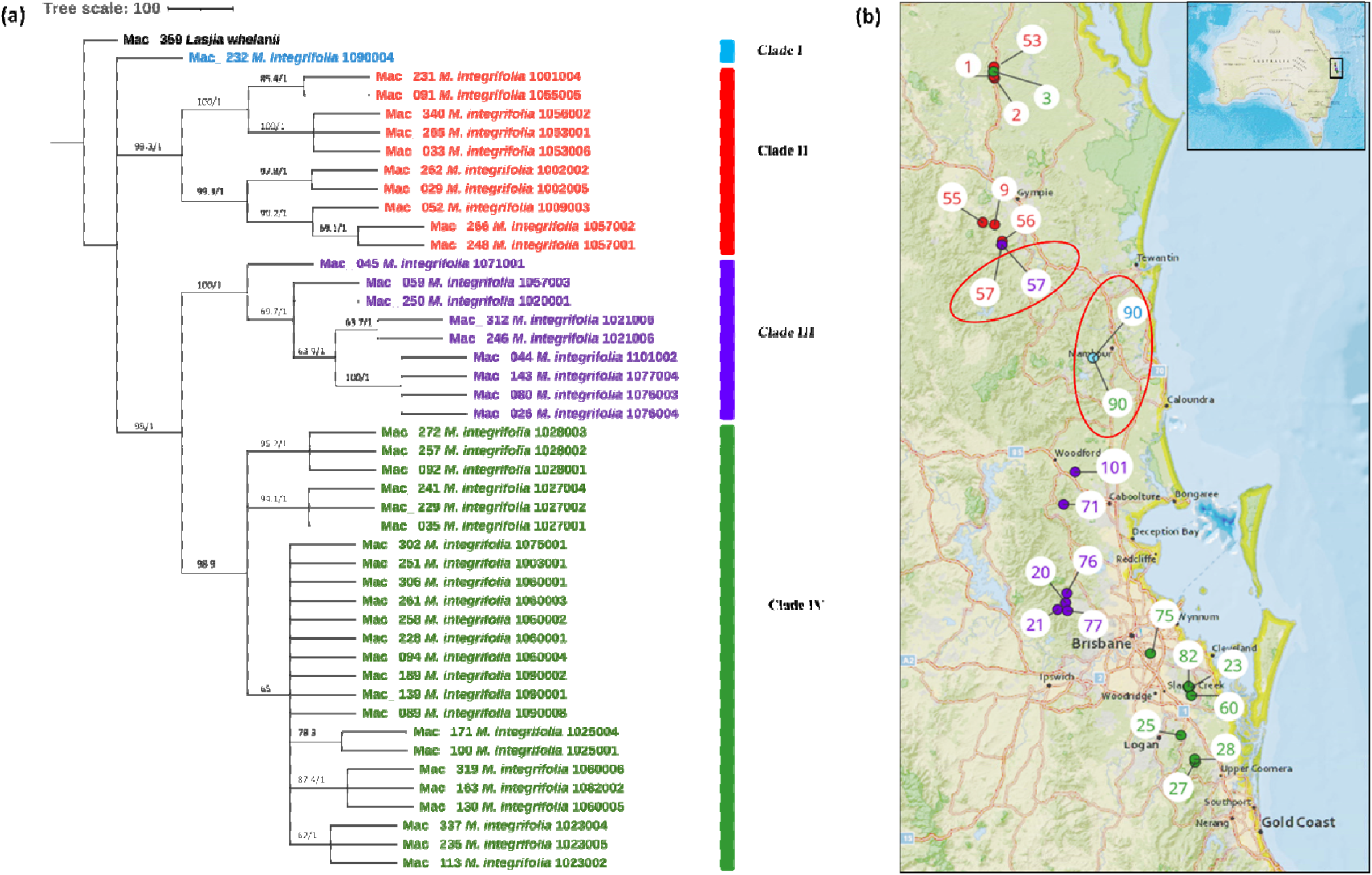
(a) Chloroplast phylogeny of *M. integrifolia* using ML and BI method. Numbers above the lines represent ML bootstrap support/Bayesian posterior probabilities. **(b) Phylogeographic analysis of *M. integrifolia.*** Numbers indicate corresponding population site (Supplementary table 4). Light blue: Clade I, Red: Clade II, Purple: Clade III and Green: Clade IV. Coloured dots on the map indicate the corresponding clade in chloroplast phylogenetic tree. Two red circles highlight population site number which contained accessions from different clades.

To study the phylogenetic relationship of *M. tetraphylla*, phylogenetic trees were constructed using 49 complete chloroplast genome sequences. The multiple chloroplast genome alignment of *M. tetraphylla* accessions together with the outgroup *L. whelanii* was 161,554 bp in length with 96% identical sites. ML and BI trees exhibited similar phylogenetic topologies (Figure 3a). The resulting phylogenetic tree showed strong statistical support for most internal and external nodes but barring poor BS value for some internal nodes. However, in Bayesian chloroplast tree, highest PP value of 1 was observed for all the nodes. Chloroplast tree displayed two major clades. The first major clade (Clade I) consisted of germplasm collected from southern part: Lismore (Mac_ 060, Mac_108, Mac_134, Mac_268, Mac_244, Mac_031, Mac_297, Mac_184, Mac_083 and Mac_095) and Ballina region (Mac_247, Mac_325, MacP_14 and Mac_115) except MacP_15 (Correspond to population site 31) and Mac_097 (Correspond to population site 38) (Figure 3b). The second major clade further divided into sub clades. Clade II contained accessions from Murwillumbah region: Mac_341, Mac_314, Mac_227, Mac_270, Mac_098, Mac_064, Mac_291, Mac_259 (Correspond to population site 37, 37, 81, 160, 160, 160, 160 and 160 respectively) and Beenleigh region; Mac_236 (Correspond to population site 100). Two accessions from Clade III from population site 37 clustered separately from rest of the accessions from same geographical location. However as in *M. integrifolia* chloroplast phylogenetic tree, the majority of *M. tetraphylla* tended to form distinct clusters among themselves based on geographical areas. Results also indicated that Mac_031, Mac_244, Mac_268 from population site 96 (Clade I) and Mac_264 and Mac_238 from population site 84 (Clade VI) were highly diverged accessions.

**Figure 3:**
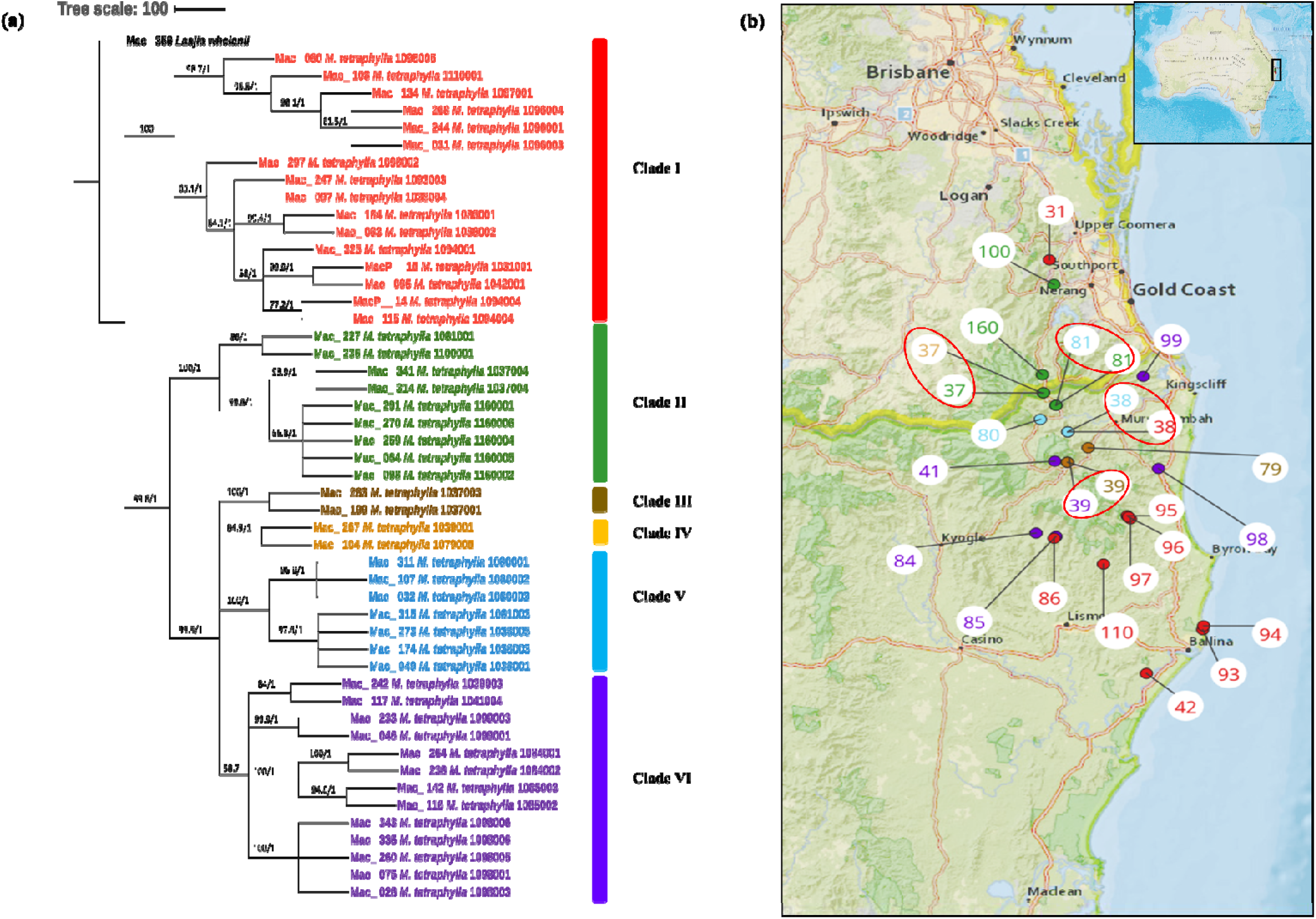
(a) Chloroplast phylogeny of *M. tetraphylla* using ML and BI method. Numbers above the lines represent ML bootstrap support/Bayesian posterior probabilities. **(b) Phylogeographic analysis of *M. tetraphylla*.** Numbers indicate corresponding population site (Supplementary table 4). Red: Clade I, Green: Clade II, Brown: Clade IIII, Orange: Clade IV, Light blue: Clade V and Purple: Clade VI. Coloured dots on the map indicate the corresponding clade in chloroplast phylogenetic tree. Four red circles highlight population site number which contained accessions from different clades.

The phylogenetic relationships within M. *ternifolia* accessions were inferred by 22 assembled complete chloroplast genomes. A multiple chloroplast alignment conducted using outgroup was 160,807 bp with 97.2 identical sites. Phylogenetic trees built with the whole chloroplast genome using both methods had the same topology (Figure 4a). The results showed two major clades having MacP_11, Mac_309 and MacP_12 from Nambour (Corresponds to population site 88) (Figure 4b) in one and remaining accessions in second major clade. There was a clear relationship between the phylogenetic structure and geographic origin of the wild accessions of *M. ternifolia*. The resulting topology suggested accessions from Clade IV: Mac_299, Mac_071, Mac_317, Mac_332 and Mac_334 (Correspond to population site 20, 51, 51, 51 and 51 respectively) were highly diverged accessions. In *M. jansenii* population no variants were observed indicating that all accessions shared the same chloroplast haplotypes except for MacP_16 with 2 bp difference.

**Figure 4:**
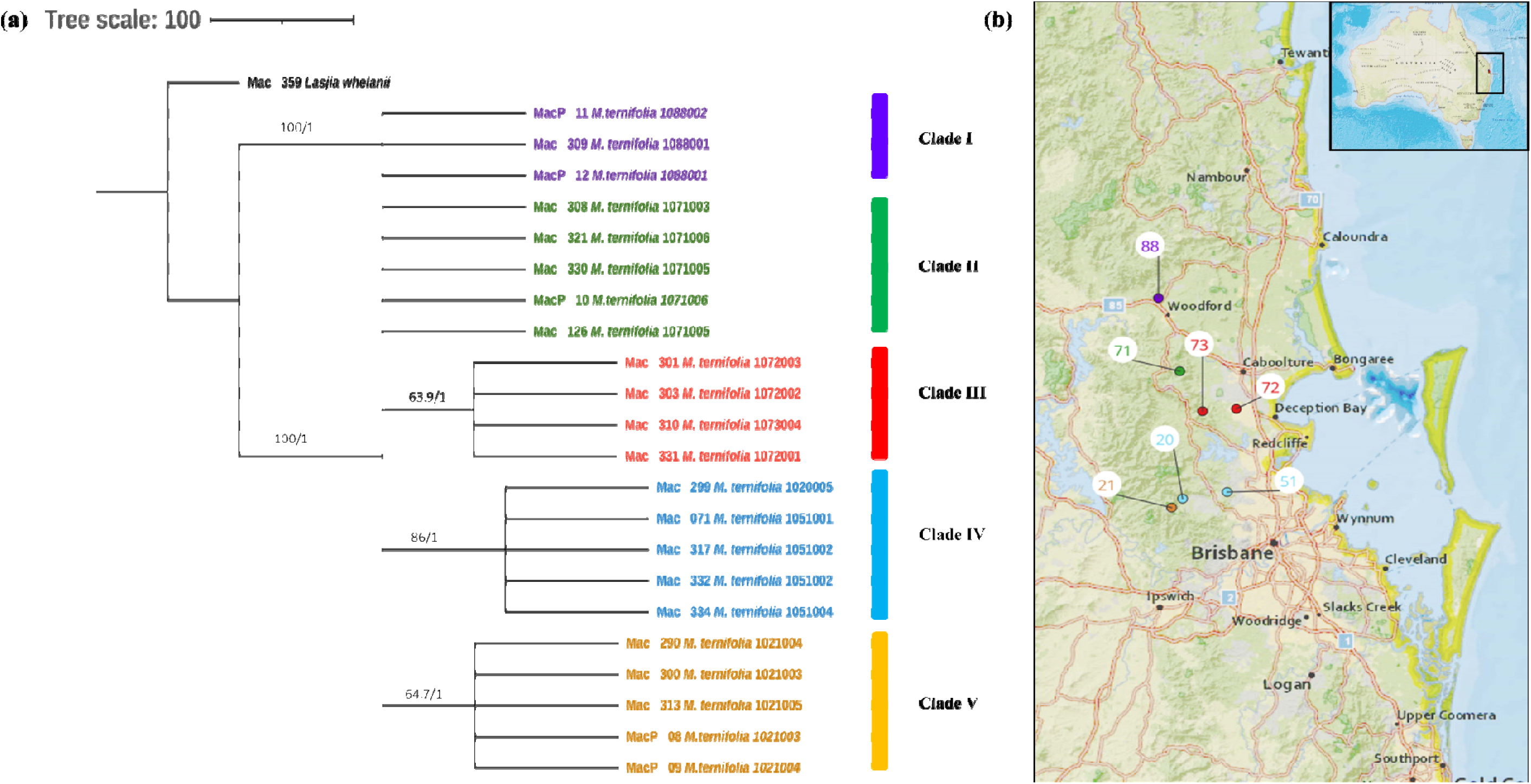
(a) Chloroplast phylogeny of *M. ternifolia* using ML and BI method. Numbers above the lines represent ML bootstrap support/Bayesian posterior probabilities. **(b) Phylogeographic analysis of *M. ternifolia*.** Numbers indicate corresponding population site (Supplementary table 4). Purple: Clade I, Green: Clade II, Red: Clade IIII, Light blue: Clade IV and Orange: Clade V. Coloured dots on the map indicate the corresponding clade in chloroplast phylogenetic tree.

Chloroplast phylogeny tree generated by taking 138 complete chloroplast genomes were supported with PP of 1.0. Two major clades were identified (Figure 5). First major clade contained 16 *M. tetraphylla* accessions from Lismore region (Mac_ 060, Mac_108, Mac_134, Mac_268, Mac_244, Mac_031, Mac_297, Mac_184, Mac_083 and Mac_095), Ballina region (Mac_247, Mac_325, MacP_14 and Mac_115), Beenleigh region (MacP_15) and Murwillumbah region (Mac_097). Interestingly all these accessions corresponded to Clade I in *M. tetraphylla* chloroplast phylogenetic tree (Figure 3a). Second major clade further differentiated into two sub clades. All the *M. jansenii* accessions were clustered in one clade. Second sub clade further divided into two clades. Small sub clade contained 10 accessions from the northern distribution of *M. integrifolia* (Correspond to Clade II in *M. integrifolia* Cp phylogenetic tree) and one accession from Nambour region (Correspond to Clade I in *M. integrifolia* Cp phylogenetic tree). Larger sub clade contained all remaining *M. tetraphylla*, *M. integrifolia* as well as *M. ternifolia.* This result shows that accessions that were collected from same locality cluster together. We assumed chloroplast capture could be the reason for the presence of different species in same clade when a species coexists in the same geographic area with other species.

**Figure 5:**
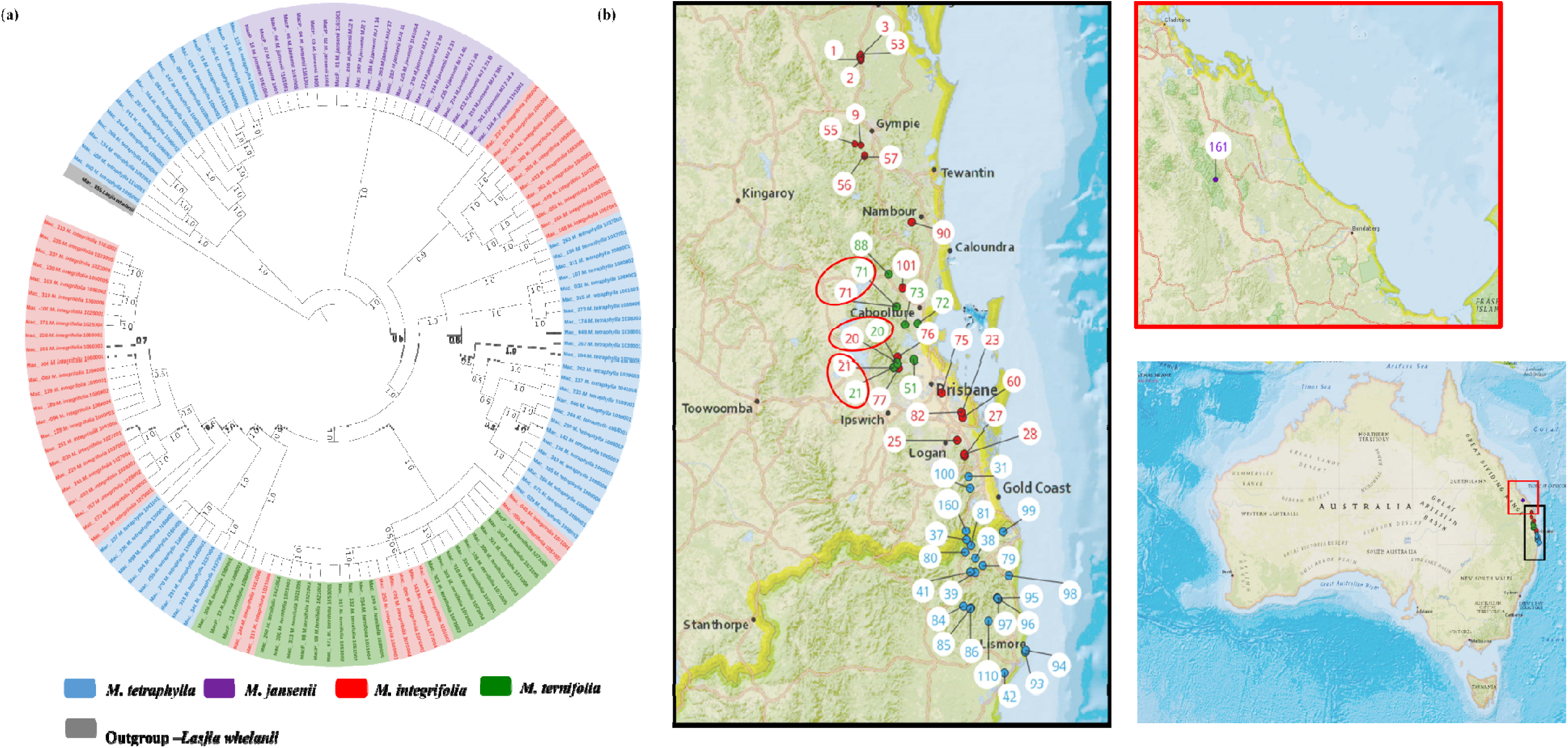
Phylogeographic results of *Macadamia* species. (a) Chloroplast phylogenetic tree of Macadamia using Bayesian inference (BI) method. Light blue: *M. tetraphylla*, Purple: *M. jansenii*, Red: *M. integrifolia* and Green: *M. ternifolia*. Numbers above the lines represent Bayesian posterior probabilities. (b) Map of Australia showing origins of Macadamia accessions. Numbers indicate corresponding population site (Supplementary table 4). Coloured dots on the map indicate the corresponding species. Three red circles highlight population site number which contained accessions from different species.

### 3.3. Nuclear gene phylogenetic analysis

For *M. integrifolia*, we used a total of 44 accessions. The multiple sequence alignment was 81,747 bp in length with 91% identical sites. The topology of nuclear gene phylogenetic tree constructed based on both ML and BI methods was nearly identical (Supplementary Figure 1 & 2). However, the resulting phylogenetic trees exhibited low bootstrap values (<70) and Bayesian posterior probabilities (<0.95). Moreover, this result was not congruent with chloroplast phylogenetic tree results.

Nucleotide alignment of 49 *M. tetraphylla* along with the outgroup was 81,753 bp in length. The phylogeny obtained with the ML approach was nearly identical to the BI approach (Supplementary Figure 3 & 4). As in *M. integrifolia* branching support rate is low. Tree topology of nuclear gene phylogeny and chloroplast phylogeny is dissimilar. Next, we constructed nuclear phylogenetic trees for *M. ternifolia* (Supplementary figure 5 & 6). and *M. jansenii* (Supplementary figure 7 & 8) population with ML method and BI method. However, resulting topologies had poor statistical support for internal and external nodes.

A ML tree was also constructed for 138 macadamia genotypes based on the concatenated single copy nuclear gene CDS. ML tree demonstrated the presence of four distinct species in Genus *Macadamia*. This well supported tree classified the population into two main clades (Figure 6). Main clade I consisted of *M. ternifolia* and *M. jansenii* while clade II consisted of all *M. integrifolia* and *M. tetraphylla*. This outcome underscored the close relationship between *M. ternifolia* and *M. jansenii* and also between *M. integrifolia* and *M. tetraphylla*.

**Figure 6:**
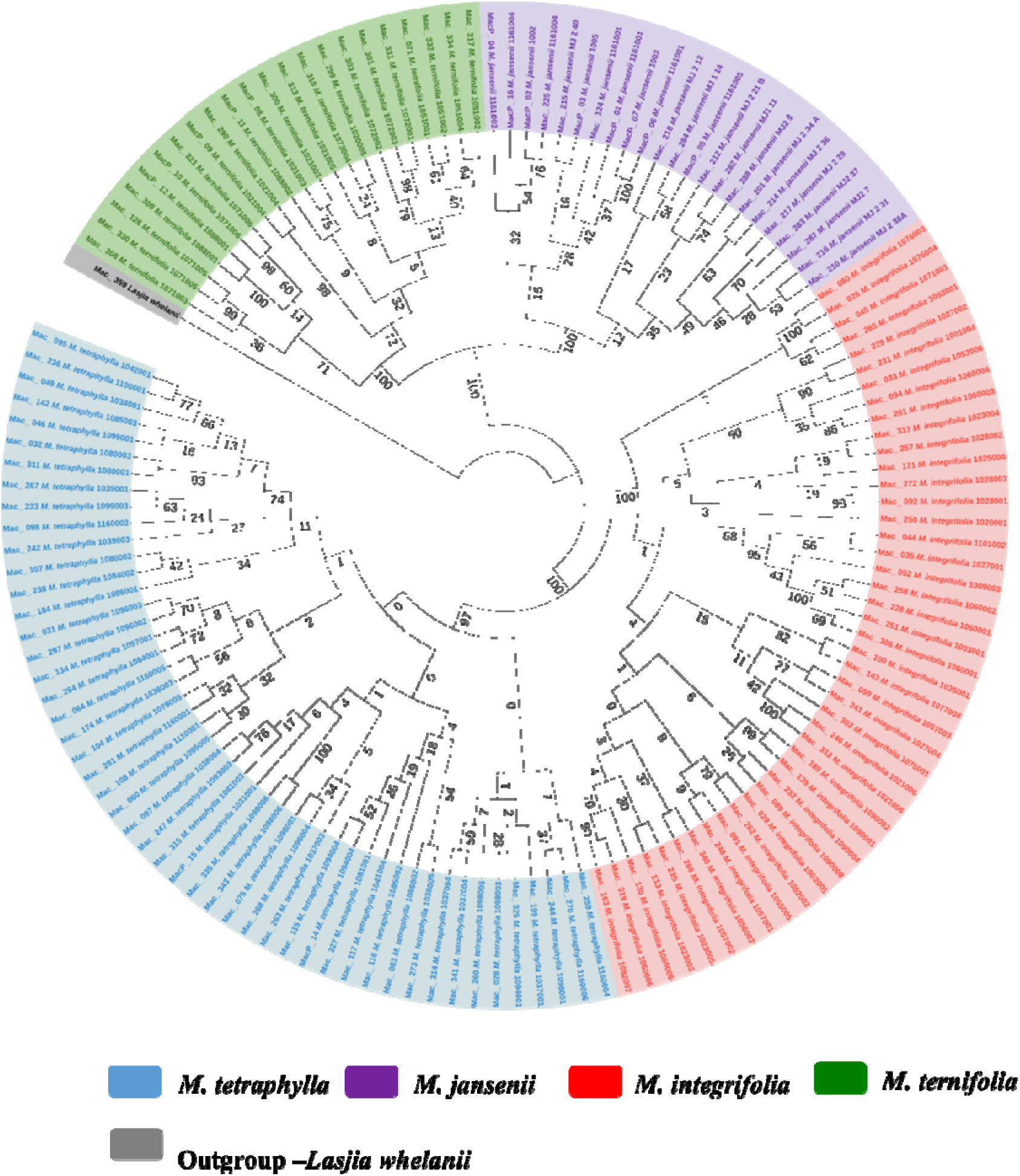
Nuclear phylogenetic results of *Macadamia* species. Light blue: *M. tetraphylla*, Purple: *M. jansenii*, Red: *M. integrifolia* and Green: *M. ternifolia*. Numbers above the lines represent ML bootstrap support. Phylogenetic tree constructed from coding sequences of 53 single copy genes using 1000 bootstrap replicates. Accessions were colour coded according to the species.

## 4. Discussion

Phylogenetics helps to unravel evolutionary histories and provides valuable insights into the factors driving the growth and adaptation of important plant groups worldwide (Lan *et al*., 2022). The current study, using whole genome of sequencing, has resolved the phylogeny of the four *Macadamia* species and confirmed that all the wild accessions belonged to 4 distinct species. Chloroplast genome sequences have been extensively used in phylogenetics analysis (Sun *et al*., 2020). Chloroplasts are the most metabolically active organelle found in plant that carry out most of the biochemical synthesis process which requires for the function of the cell in producing energy through photosynthesis (Dobrogojski *et al*., 2020). The chloroplast DNA sequence has unique features compared to nuclear genome in analysis for population genetics and evolutionary relationships within families, genus, and species (Ahmad Termizi *et al*., 2016; De Las Rivas *et al*., 2002). The chloroplast genome is present in high copy numbers, has a low rate of spontaneous mutation, and does not undergo cross overs or recombination. Chloroplast genome sequence data are highly conserved (Dobrogojski *et al*., 2020). Earlier research relies on separation of chloroplast genome from the nuclear genome and mitochondrial genome (Jansen *et al*., 2005). With the emergence of NGS technology new high throughput approaches have been introduced for successful isolation of chloroplast sequencing with low cost (Ahmad Termizi *et al*., 2016). The read length, sequencing depth, sequence coverage or width, and evenness of coverage influence accuracy of a DNA sequencing using NGS technology. Short read sequencing has been successfully applied to sequence chloroplast genomes of various plant species (Kyriakidou *et al*., 2018). Although there were numerous records of extensive use of chloroplast genomes in evolutionary relationship in plants, very few studies were presented on whole chloroplast genome data in macadamia. In this study, sizes of all chloroplast genomes were consistent with previous macadamia chloroplast genomes (Liu *et al*., 2017; Liu *et al*., 2018; Nock *et al*., 2014). Genomes annotation resulted in higher number of genes compared to previous studies (Liu *et al*., 2017; Liu *et al*., 2018; Nock *et al*., 2014) which reported 79 coding sequences, 4 rRNA and 30 tRNA for *M. integrifolia* (Nock *et al*., 2014), *M. ternifolia* (Liu *et al*., 2017) and *M. tetraphylla* (Liu *et al*., 2018). The difference in the gene number was possibly due to the difference in the annotation tool. All the previous genomes were annotated using Dual Organelle GenoMe Annotator (DOGMA) (Wyman *et al*., 2004) while the current genomes were annotated by GeSeq online tool (https://chlorobox.mpimp-golm.mpg.de/geseq.html). Moreover, there is no previously reported chloroplast genome for *M. jansenii*. For the first time, we have generated chloroplast genomes for 23 *M. jansenii* accessions using the Get Organelle toolkit (Jin *et al*., 2020).

This study provides the most comprehensive analysis of the evolutionary relationships of the chloroplasts within the species in the genus *Macadamia*. The topology of the chloroplast phylogenetic tree with distinct northern population and southern population of *M. integrifolia* is in agreement with the previously published phylogenetic results (Nock *et al*., 2019) (Lin *et al*., 2022). Nock *et al*. (Nock *et al*., 2019) also reported two distinct populations namely Gundiah/Mount Bauple population and Gympie population in the northern clade. A similar result was also reported by Lin, J., *et al*. (2022) (Lin *et al*., 2022) based on chloroplast and nuclear phylogenetic analysis. However, our results do not clearly separate accessions between Gundiah/Mount Bauple region: Mac_231, Mac_262, Mac_029, Mac_265 and Mac_033 (Corresponding to population sites 1, 2, 2, 3 and 3 respectively) (Figure 2b) and Gympie region: Mac_052, Mac_091, Mac_340, Mac_248 and Mac_266 (Corresponding to population sites 9, 55, 56, 57 and 57, respectively). In this study, the ML/BI phylogenetic tree showed that Mac_232 from Clade I (Figure 2b) was clustered separately from other three accessions in Cluster III (Mac_089, Mac_139 and Mac_189) originating from population site 90, suggesting that it is a planted tree. This finding was also supported by the previous SSR results, which indicated that the Dulong tree is a planted tree which originated from the Brisbane region (Nock, 2022). The results also show that Mac_059 from the population site 57 is a planted tree, where this was not reported previously. Moreover, all the trees from Sunshine coast: Mac_089, Mac_139, Mac_189 (Figure 2a-Clade IV) are likely to be planted trees which is consistent with previous studies (Nock, 2022; Nock *et al*., 2019). The results revealed that Mac_026, Mac_044, Mac_080 and Mac_143 are highly diverged accessions, in contrast to the previous study (Mai *et al*., 2020), in which Mac_229, Mac_266, Mac_139 and Mac_235 were recognized as diverged accessions. The phylogeographic results of the present study were in agreement with those of the previous study by Mai *et al*. (2020) (Mai *et al*., 2020). Chloroplast phylogenetic analysis of *M. tetraphylla* revealed that MacP_15 (Correspond to population site 31), Mac_097 (Correspond to population site 38) and Mac_236 (Correspond to population site 100) (Figure 3a & 3b) might have been moved by humans as they were clustered with accessions from a different locality. It is noteworthy that we identified MacP_15 and Mac_236 were outside the range of the natural population. *M. tetraphylla* is mostly distributed in the New South Wales region (Topp *et al*., 2019). There is no record of natural occurrence in the Beenleigh, QLD region. Previous studies reported a weak genetic differentiation (Mai *et al*., 2020; Nock, 2022; O’Connor *et al*., 2015) for *M. tetraphylla* populations. However, the present study revealed a positive correlation between genetics and geographical distribution.

For the first time, we report the phylogeographic pattern of distribution of genetic variation for the *M. ternifolia* population. However, further investigation is needed with an increased number of samples. Results for the *M. jansenii* accessions identified the presence of one chloroplast haplotype as expected due to the small, isolated population. This suggested that *M. jansenii* has gone through a genetic bottleneck. *M. jansenii* is found only in Bulburin National Park north of Bundaberg that is 180 km away from any *M. integrifoli*a population (Mai *et al*., 2020; Topp *et al*., 2019). Therefore, the possibility of gene flow between the two population is limited except for the movement of nuts with the involvement of human. A decrease in the movement of genes is expected to increase the occurrence of inbreeding among the individuals in the population (Hatmaker *et al*., 2018). Inbreeding, in turn can have effects on the genetic health of the population, potentially leading to an accumulation of harmful traits and decrease in overall fitness (Hatmaker *et al*., 2018).

In contrast to the previously recorded phylogenies (Mai *et al*., 2020; Mast *et al*., 2008; Nock, 2022; Peace, 2005), we found that accessions that were collected from the same geographical location were closely related. The distinct separation of *Macadamia* populations within species reported in previous phylogenies (Mai *et al*., 2020; Mast *et al*., 2008; Nock, 2022; Peace, 2005) were based upon chloroplast genome analysis. This study shows that reticulate evolution has resulted in chloroplast transfer between species and resulted in distinct chloroplast types within individual species but has not impacted on the distinctness of the nuclear genomes. Although chloroplast capture was not previously reported in Macadamia, many other plants have reported the occurrence of reticulate evolution of the chloroplast (Acosta and Premoli, 2010; Ananda *et al*., 2021; Moner *et al*., 2020; Wambugu *et al*., 2015; Yi *et al*., 2015). In this study, chloroplast phylogeny separated 16 *M. tetraphylla* accessions from the rest of the accessions. The second major clade further divided into two clades having all *M. jansenii* in one clade and rest of the accessions in the other. Furthermore, a sub clade further separated 11 *M. integrifolia* and 24 *M. tetraphylla* accessions leaving a complex clade having *M. integrifolia*, *M. ternifolia* and *M. tetraphylla*. This suggested a series of chloroplast capture events between *M. integrifolia*, *M. ternifolia* and *M. tetraphylla*.

Phylogenetic trees built with the single copy nuclear gene CDS for *Macadamia* species strongly supported four distinct species in the genus *Macadamia* as reported in previous studies (Nock, 2022; Peace, 2005). Individual nuclear phylogenetic trees for the four species showed little structure, suggesting widespread gene flow within each species and little geographic structure in the nuclear genome.

The availability of large sequence data has significantly advanced our understanding of the *Macadamia* species distribution and diversity essential for both conservation and breeding programs. This advanced knowledge aids in the conservation of these species, now found in fragmented rainforest habitats by highlighting the importance of in situ conservation strategies that focus on capturing a wide range of genetic diversity within sites. Such conservation efforts are crucial for safeguarding the species against extinction but also in enhancing their commercial value and sustainability for future generation.

## Acknowledgements

We gratefully acknowledge University of Queensland Research Computing Centre (UQ-RCC) for providing all the computation resources. We would like to thank Dr. Catherine J Nock for providing helpful comments on phylogenetic analysis. We thank Denise Bond from Macadamia Conservation trust for generously sharing data.

## Supplementary data

Supplementary figure 1: Nuclear gene phylogeny of *M. integrifolia* using Maximum likelihood (ML) method.

Supplementary figure 2: Nuclear gene phylogeny of *M. integrifolia* using Bayesian inference (BI) method.

Supplementary figure 3: Nuclear gene phylogeny of *M. tetraphylla* using Maximum likelihood (ML) method.

Supplementary figure 4: Nuclear gene phylogeny of *M. tetraphylla* using Bayesian inference (BI) method.

Supplementary figure 5: Nuclear gene phylogeny of *M. ternifolia* using Maximum likelihood (ML) method.

Supplementary figure 6: Nuclear gene phylogeny of *M. ternifolia* using Bayesian inference (BI) method.

Supplementary figure 7: Nuclear gene phylogeny of *M. jansenii* using Maximum likelihood (ML) method.

Supplementary figure 8: Nuclear gene phylogeny of *M. jansenii* using Bayesian inference (BI) method.

Supplementary Table 1. Details of the samples used in the study

Supplementary Table 2. Details of raw and trimmed data

Supplementary Table 3. Details of selected 53 single copy nuclear gene CDS in *M. integrifolia* (GCF 013358625.1)

Supplementary Table 4. Details of 138 wild accessions using in phylogenetic study.

Supplementary Table 5. Details of chloroplast annotation of 166 macadamia accessions and *Lasjia whelanii*

Supplementary Table 6. Details of chloroplast genes identified in macadamia genome

Supplementary Table 7. NCBI submission details of 166 wild accessions and Lasjia *whelanii* using in the study.

## Author contribution

**Robert J. Henry:** Design, Methodology, Data collection, Supervision, Project administration, Funding acquisition, Resources and Writing - Review & Editing. **Agnelo Furtado:** Design, Methodology, Data collection, Supervision, Project administration, Formal analysis, Supervision, Resources, Data Curation, Writing - Review & Editing. **Bruce Topp:** Design, Methodology, Data collection, Supervision, Data Curation, Writing - Review & Editing. **Mobashwer Alam:** Methodology, Data collection, Supervision, Writing - Review & Editing**. Patrick J. Mason:** Methodology, Data collection, Writing - Review & Editing. **Ardashir Kharabian-Masouleh:** Methodology, Formal analysis, Software, Writing - Review & Editing. **Sachini Lakmini Manatunga:** Formal analysis, Data curation, Writing original draft.

## Data Availability

All sequence data will be available at NCBI via Bio Project: PRJNA1036028 with Bio sample number SAMN38356272 - SAMN38356438 (Supplementary table 7).

